# Evaluating a Cassava Crop Growth Model by Optimizing Genotypic-Specific Parameters Using Multi-environment Trial Breeding Data

**DOI:** 10.1101/2024.10.29.620843

**Authors:** Pamelas M. Okoma, Siraj S. Kayondo, Ismail Y. Rabbi, Patricia L. Moreno-Cadena, Gerrit Hoogenboom, Jean-Luc Jannink

## Abstract

Cassava (*Manihot esculenta* Crantz) is a critical food security crop for sub-Saharan Africa. Efforts to improve cassava through breeding have expanded over the past decade. At the same time, crop growth models (CGM) are becoming common place in breeding efforts to expand the inference of evaluations of breeding germplasm to environments that have not been tested and to prepare for breeding for adaptation to future climates. We parameterized a CGM, the CROPGRO-MANIHOT-Cassava model in the DSSAT family of models, using data on 67 clones from the International Institute of Tropical Agriculture cassava breeding program evaluated from 2017 to 2020 and over eight locations in Nigeria using trial and error parameter adjustments and the General Likelihood Uncertainty Estimation method. Our objectives were to assess the feasibility of this large-scale calibration in the context of a cassava breeding program and to identify systematic biases of the model. For each cultivar we calculated the Pearson correlation between model prediction and observation across the environments, as well as root mean squared error and d statistics. As a result of calibration, the correlation coefficient increased from −0.03 to +0.08, the RMSE dropped from 21 t ha^-1^ to 5 t ha^-1^ while d increased from 0.23 to 0.44. We found that the model underestimated root yield in dry environments (low precipitation and high temperature) and overestimated root yield in wet environments (high precipitation and low temperature). Our experience suggests both that CGM calibration could become a routine component of the cassava breeding data analysis cycle and that there are opportunities for model improvement.

## Introduction

Crop Growth Models (CGMs) now have a long history in the study of agronomic management practices (Jones et al. 2003; Holzworth et al. 2018; Jones et al. 2017; Tsuji et al. 1998). CGMs simulate plant phenology and growth by integrating ecophysiological functions with weather data, usually on a daily time step, tracking photosynthesis and partitioning as well as resource extraction from the environment. Once calibrated, a CGM can be used to study such questions as planting dates or fertilizer application (Banterng et al. 2009), conservation tillage (Corbeels et al. 2016), crop rotations (Mohanty et al. 2012), or even intercropping (Berghuijs et al. 2021). An advantage of studying such issues with a model is that the system response can be predicted over a long timeframe or for many different location conditions representing a breadth of environments that would be impossible to evaluate empirically. Even for well-calibrated models, results cannot be taken at face value but should suggest the important factors to explore with field experiments.

Expanding the use of CGMs from agronomic questions to plant breeding has been more recent (Baenziger et al. 2004; Technow et al. 2015; Messina et al. 2018; White et al. 1996). Plant breeders base selection decisions on predictions of future performance of breeding lines, whether those predictions come from prior observations of the lines themselves or of close relatives whose genetic relatedness can be inferred from DNA variants. In either case, the predictions cannot be extrapolated to environments or management practices that have not been observed.

Thus, a motivation to parameterize CGMs for breeding lines would be to enable predictions in environments where they have not been tested. Applications of such predictions could be estimating the stability of a line’s performance by modeling outcomes over many years or locations (for example, aiding such efforts as Lozada and Carter 2020; Olivoto et al. 2021), identifying specific environments that may be most promising for specific breeding lines (as Chai, et al. 2022 have suggested occurs in modern breeding, for example, Phoncharoen, et al. 2021) or for identifying optimal locations for a multi-location trial evaluation network (Putto et al. 2008). These models may therefore contribute to the development of efficient breeding strategies through decision support in breeding programs. A number of these applications are reviewed in the context of breeding to adapt to climate change in (Ramirez-Villegas et al. 2020).

As for many developments in breeding, attempts to integrate CGMs into breeding have been spurred on by the plummeting cost of DNA markers that has made it possible to associate traits with alleles or haplotypes at specific genomic segments. A challenge to the use of CGMs in breeding is the effort and cost to estimate many CGM parameters for individual breeding lines, as exemplified by early efforts (e.g., White and Hoogenboom 1996; Hoogenboom and White 2003). In particular, CGMs require many parameters that are not routinely measured in breeding programs so that they come at an extra cost to the program, which is prohibitive. A work around to this problem is to seek to estimate parameter values from data that *are* routinely collected in breeding programs. Estimation procedures essentially fit parameter values so that CGM predictions match as closely as possible to observed breeding data. One such method is General Likelihood Uncertainty Estimation (GLUE) that has been incorporated in the Decision Support System for Agrotechnology Transfer (DSSAT) (He et al. 2010; Hoogenboom et al. 2021; Ferreira et al., 2024). To extend this methodology to cassava (Manihot esculenta Crantz), the initial step involves the development of a robust growth model.

The worldwide importance of cassava is undeniable, especially in sub-Saharan Africa where cassava roots are a valuable source of calories (Adebayo 2023; Chiaka et al. 2022). Numerous efforts to improve cassava breeding to deliver greater genetic gain are underway. Such efforts span a wide range of technologies from on farm testing (Nanyonjo et al. 2024), to genomic selection (Wolfe et al. 2017; Ceballos et al. 2021), to developing cassava hybrids (Zhang et al. 2024). Integrating crop growth models into these efforts could prove beneficial for optimizing field evaluations, matching variety candidates to target environments, as well as for experimenting with new management options. A number of cassava CGMs have been reviewed (Phoncharoen, et al. 2021, Moreno-Cadena et al. 2021). An advantage of the CSM-MANIHOT-Cassava model described there is that is has been integrated within the DSSAT Crop Modeling Ecosystem in DSSAT v. 4.8 (Hoogenboom et al. 2019) for which there is also an R interface (Alderman 2020). The DSSAT system in turn incorporates model calibration functionality, including GLUE.

The MANIHOT-Cassava model was developed from the CSM-CROPSIM model, itself a development of the GUMCAS model (Moreno-Cadena et al. 2020). While (Moreno-Cadena et al. 2020) only verbally describes the model updates, comprehensive details including equations are available in (Moreno-Cadena 2021). Plant development is a function of accumulated thermal time with different cardinal temperatures for the processes of forking and leaf growth. Daily photosynthetic assimilates are obtained by multiplying the solar radiation intercepted by the photosynthetically active radiation use efficiency (PARUE). The model uses a spill-over strategy for biomass allocation: assimilates are allocated by prioritizing the growth of aboveground biomass and fibrous roots according to their demand and the remainder is assigned to the storage roots. Drought stress dynamics are influenced by soil moisture content, impacting germination, branching, leaf appearance, and photosynthesis. Nitrogen limitations are also factored in, though they necessitate further refinement. The current model version requires an array of parameters at the species, ecotype, and cultivar levels, which define stress thresholds, senescence processes, Vapor Pressure Deficit (VPD) effects, root water uptake, and other physiological traits like leaf morphology and growth characteristics. As a DSSAT crop growth model, MANIHOT-Cassava requires minimum data sets including daily weather data, soil surface and profile characteristics, crop management, initial conditions, and crop measurements (Hoogenboom et al. 2012, 2019).

The model performed well in simulating cassava development and growth, including forking dates, leaves and stem biomass, and the yield and yield components (Moreno-Cadena et al. 2020; Phoncharoen, et al. 2021). A previous study assessed the MANIHOT-Cassava model as a breeding tool for identifying the most suitable production environments for potential varieties in Thailand (Phoncharoen, et al. 2021). Nevertheless, any model is a work in progress with the potential to better track responses of many growth processes to external conditions and attempting to use more parsimonious parameterizations. Our objectives were to 1) assess the feasibility of calibrating the model based on observations made by the International Institute of

Tropical Agriculture (IITA) breeding program using GLUE and across a range of elite Nigerian cassava germplasm and growing environments and 2) to identify any systematic biases in model predictions after that calibration, to suggest further improvements to the model. The experimental data included 67 clones evaluated across 11 locations in three years, thus representing a large dataset for model assessment. This report presents the first calibration of the CSM-MANIHOT-Cassava model in Nigeria, a country with a wide range of environmental conditions for cassava cultivation.

## Materials and Methods

### Model calibration

#### Experimental design and fields management

Phenotypic data were collected for 67 clones from the Uniform Yield Trials of IITA, which are the most advanced evaluation stage of the cassava breeding program prior to variety release decisions. The clones were derived from crosses within IITA’s elite population as part of the genomic recurrent breeding program. They underwent four stages of screening for diseases, vigor, agronomic, and quality traits in earlier trials of the breeding program, showing resistance to diseases and potential for high yield and dry matter content. To standardize the model terminology, we call clone a “cultivar”.

The trials were targeted across 11 IITA testing stations in Nigeria (Figure 1) over three growing seasons (2017-2018, 2018-2019, and 2019-2020). We considered each year by location combination to represent one environment. The fields selected for the trials had been fallow, with no crops grown in the previous three years. Before planting, the selected field were mowed, ploughed, harrowed, and ridged. The planting date varied from April to August depending on the rainy season in each location (Table 1). Cultivars were planted in a randomized complete block design with 3 replications. Each plot consisted of 6 rows of 5.6 m (seven plants) with an inter-row spacing of 1 m and intra-row spacing of 0.8 m (plot size = 28 m^2^). The trials were managed without pesticides, irrigation, or fertilizer.

**Table 1.** Description of experimental sites in Nigeria.

| Location | Lat / Long | Soil | YEAR | Name | Planting Date | TMIN<br>(°C) | TMAX<br>(°C) | RAIN<br>(mm) | RH2M<br>(%) | SRAD<br>(MJ/m<sup>2</sup>/day) | WIND<br>(m/s) |
| --- | --- | --- | --- | --- | --- | --- | --- | --- | --- | --- | --- |
| ABUJA | 9.07°N, 7.47°E | SCL | 2019-2020 | Abuja19 | June 2019 | 20.8 | 30.4 | 2238 | 73.2 | 18.6 | 1.6 |
| AGO-OWU | 7.26°N, 4.32°E | SCL | 2017-2018 | Ago17 | August 2017 | 21.8 | 29.2 | 1479 | 85.9 | 16.7 | 1 |
| AGO-OWU | 7.26°N, 4.32°E | SCL | 2018-2019 | Ago18 | June 2018 | 22.0 | 28.9 | 2842 | 87.1 | 16.6 | 0.9 |
| AGO-OWU | 7.26°N, 4.32°E | SCL | 2019-2020 | Ago19 | June 2019 | 21.7 | 29.1 | 3152 | 85.6 | 16.7 | 1 |
| IBADAN | 7.47°N, 3.93°E | SL | 2017-2018 | Ibadan17 | September 2017 | 22.0 | 29.1 | 2179 | 86.4 | 16.5 | 1.5 |
| IBADAN | 7.47°N, 3.93°E | SL | 2018-2019 | Ibadan18 | July 2018 | 22.2 | 29.3 | 2316 | 86.5 | 16.9 | 1.4 |
| IBADAN | 7.47°N, 3.93°E | SL | 2019-2020 | Ibadan19 | May 2019 | 21.8 | 29.4 | 2397 | 85 | 16.6 | 1.5 |
| IKENNE | 6.87°N, 3.71°E | CL | 2017-2018 | Ikenne17 | August 2017 | 22.8 | 29.2 | 2075 | 88.2 | 16.3 | 1.1 |
| IKENNE | 6.87°N, 3.71°E | CL | 2018-2019 | Ikenne18 | June 2018 | 23.0 | 29.3 | 2533 | 88.4 | 16.5 | 1.1 |
| IKENNE | 6.87°N, 3.71°E | CL | 2019-2020 | Ikenne19 | June 2019 | 22.9 | 29.3 | 3038 | 88 | 16.5 | 1.1 |
| MOKWA | 9.31°N, 5.07°E | SCL | 2017-2018 | Mokwa17 | August 2017 | 22.2 | 32.6 | 1594 | 67.8 | 18.3 | 1.7 |
| MOKWA | 9.31°N, 5.07°E | SCL | 2018-2019 | Mokwa18 | January 2018 | 22.1 | 32.4 | 1351 | 68.5 | 18.6 | 1.7 |
| MOKWA | 9.31°N, 5.07°E | SCL | 2019-2020 | Mokwa19 | June 19 | 21.8 | 32.5 | 1385 | 69.1 | 18.8 | 1.8 |
| ONNE | 4.79°N, 7.17°E | SCL | 2018-2019 | Onne18 | April 2018 | 23.3 | 29.6 | 3319 | 88.1 | 14.9 | 0.9 |
| UBIAJA | 6.67°N, 6.39°E | SCL | 2019-2020 | Ubiaja19 | June 2019 | 22.1 | 29.3 | 2725 | 84.1 | 16.7 | 1.6 |
| UMUDIKE | 5.47°N, 7.55°E | SCL | 2018-2019 | Umudike18 | July 2018 | 22.9 | 29.4 | 1966 | 86.3 | 16.2 | 1.4 |

\* Geo Loc = geographical location, SCL = Sandy Clay Loam, SL = Sandy Loam, CL= Clay Loam, Name = Location/year, TMAX = Daily temperature maximum, TMIN = Daily temperature minimum, RAIN = Sum precipitation /year, RH2M = Relative Humidity at 2 m, SRAD = Daily solar radiation, PAR = Daily photosynthetic radiation, WIND = Wind Speed at 2 m. The meteorological data corresponds to the growing season.

**Table 1:** Definition of the CSM- MANIHOT-Cassava genotype parameters in DSSAT.

| Level | Parameter | Description | Function |
| --- | --- | --- | --- |
| Cultivar | B01ND | Duration from planting to first forking (°Cd) | Phenology |
|  | B12ND | Duration from first to second forking (°Cd) | Phenology |
|  | B23ND | Duration from second to third forking (°Cd) | Phenology |
|  | B34ND | Duration from second to third forking (°Cd) | Phenology |
|  | BR1FX | Branch number per fork at fork 1 (no.) | Growth |
|  | BR2FX | Branch number per fork at fork 2 (no.) | Growth |
|  | BR3FX | Branch number per fork at fork 3 (no.) | Growth |
|  | BR4FX | Branch number per fork at fork 4 (no.) | Growth |
|  | LAXS | Maximum leaf area when growing without stress (cm<sup>2</sup>) | Growth |
|  | SLAS | Specific leaf lamina area when growing without stress (cm<sup>2</sup> g<sup>-1</sup>) | Growth |
|  | LLIFA | Leaf life from full expansion to start senescence (°Cd) | Growth |
|  | LPEFR | Leaf-petiole fraction (fraction of lamina petiole) | Growth |
|  | LNSLP | Slope for leaf production | Growth |
|  | NODWT | Node weight for the first stem of the shoot before branching at 3400 °Cd (g) | Growth |
|  | NODLT | Mean internode length (cm) for the first stem of the shoot before branching when is lignified (cm) | Growth |
| Ecotype | PARUE | Photosynthetically active radiation (PAR) conversion factor standard (g dry matter MJ<sup>-1</sup>) | Ecotype |
|  | KCAN | Photosynthetically active radiation extinction parameter (no.) | Ecotype |

**Figure 1.**
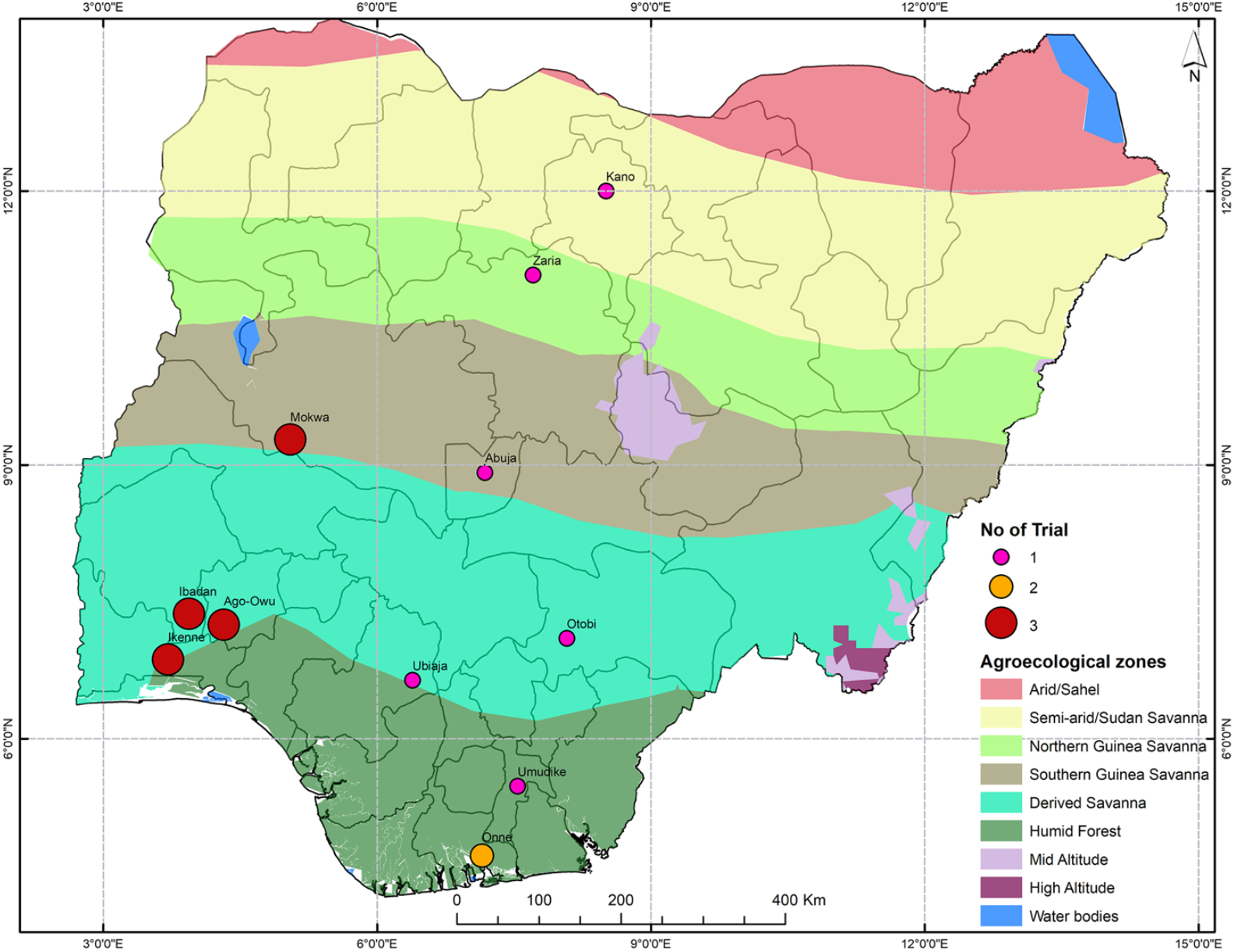
Map of Nigeria showing the locations of multi-environment trials in the IITA cassava breeding program. The agrometeorological zones and the number of tests carried out specified by the colors and the size of the point respectively.

Preliminary analysis of each trial was conducted to determine the relative strength of the genetic signal its data. The analysis was done as described by (Bakare et al. 2022) who worked on the same data. A linear mixed-effects model was fitted using the lmer function from the lme4 package in R 4.1.0 (R Core Team 2022). The model formula was:

Trait ∼ PropHar + (1|repInTrial) + (1|blockInRep) + (1|germplasmName)

- Trait = fresh storage root weight or fresh above ground biomass as responses variables,
- PropHar = proportion of plants harvested per plot (fixed effect),
- repInTrial = number of replications nested within trial (random effect),
- blockInRep = block nested within replication (random effect),
- germplasmName = cultivar’s name (random effect).

The reliability of the model was extracted using the r.squaredGLMM function from MuMIn package in R. The broad-sense heritability and the reliability for both traits (fresh storage root weight or fresh above ground biomass) were calculated by trial. Trials were filtered to retain those with H^2^ over 0.25 and reliability over 40% for at least one trait (Supplementary Figure S1). A total of 28 trials out of 38 trials were retained after filtering, included eight locations and all three years. These trials spanned 16 environments.

### Soil and weather

For each location, specific soil and weather data were used. Soil profile characteristics were obtained from the Harvard Dataverse soils data Repository for crops (Han et al. 2019) according to the GPS coordinates for each location. The soil ID code and properties are given in Supplementary Table S1. The considered parameters included: Soil Lower limit of plant extractable soil water (SLLL), Soil Drained upper limit (SDUL), Saturated upper limit (SSAT), Soil root growth factor (SRGF), Saturated hydraulic conductivity (SSKS), Soil bulk density (SBDM g/cm^3^), Soil organic carbon (SLOC), Soil clay (SLCL), Soil silt (SLSI), Soil total nitrogen (SLNI), Soil pH in water (SLHW) and Soil cation exchange capacity (SCEC).

Daily weather data were collected from the National Aeronautics and Space Administration Prediction of Worldwide Energy Resource project (https://power.larc.nasa.gov/data-access-viewer/). The data set for the growing season, summarized in the Table 1, included: minimum temperature (TMIN), maximum temperature (TMAX), precipitation (RAIN), relative humidity (RH2M), solar radiation (SRAD) and wind speed (WIND).

The DSSAT Weatherman tool was used to create input weather data files in the model format (Hoogenboom et al. 2019). The initial soil water conditions for each layer were defined by setting available soil water at 100%, the incorporation depth for surface residue at 20 cm and the simulation was done under non-nitrogen-limited and non-phosphorus-limited production conditions.

### Plant phenology

Plant phenological and leaf area index (LAI) data were collected only Ibadan in 2021. Five plants per plot, with two plots per cultivar, were assessed at 5 and 6 months after planting as follows. From the first branching level to the top of the plant, one branch was selected to be monitored at each branching point. Plant height at first branching, full plant height (from the bottom to the last leaf), length, and the number of internodes of each section between 2 forking points were measured. Three weeks later, the height and the number of internodes of new growth were measured. From these measurements, the phyllochron or time for node development was calculated. The forking time was estimated as a function of the section height and internode number. In the event of a genotype not showing forks during data collection, the default values provided by the model were used (Moreno-Cadena et al. 2020). The details of these default values can be found at: https://github.com/DSSAT/dssat-csm-os/tree/develop/Data/Genotype.

### Leaf area index

We established a predictor model to estimate leaf area. We collected fully expanded, photosynthetically active, and undamaged leaves from the top, middle, and bottom of a plant not used for phenology. Thirty leaves were sampled per cultivar. The number of lobes, length, and width of the central lobe were measured for each leaf. The leaf was scanned to calculate its area with ImageJ software bundled with Java 8 on 10/29/24 9:51:00 AM. With these data, we established a model (Equation 1) that used the length and width of the central lobe and the number of lobes to estimate the leaf area of cassava (Figure 2).

**Figure 2.**
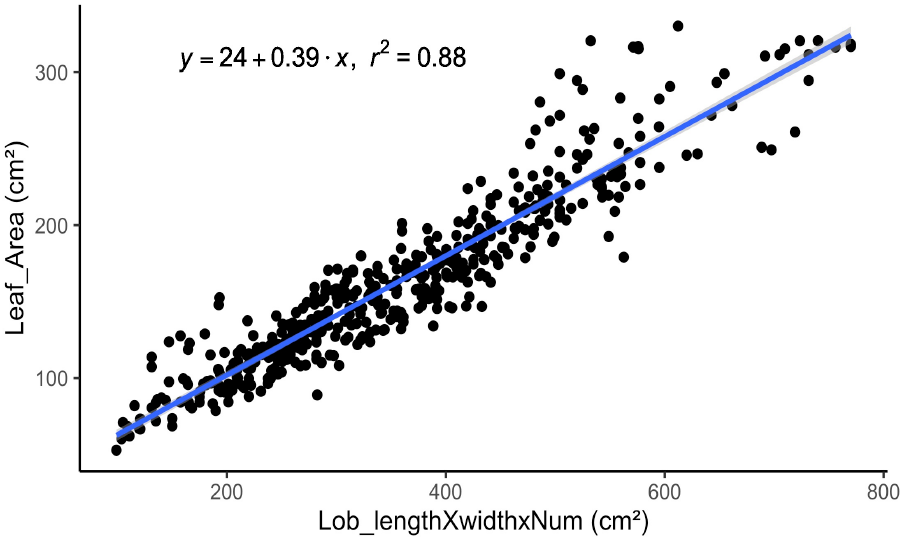
Linear regression between the product of leaf lobe length, leaf lobe width, leaf lobes number width and leaf area for 67 cassava cultivars

From the targeted plant (5 plants per plot), three leaves per plant (from top to bottom) were evaluated by the number of lobes, the length, and the width of the central lobe. The leaf area was estimated with the established equation (Equation 1). Then LAI was calculated as the total number of green leaves per plant multiplied by the mean per plant of leaf area and divided by the area occupied of each plant (Equation 2).

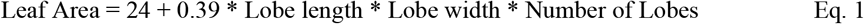

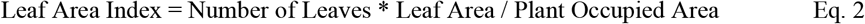

Yield and yield components were collected from 2017 to 2020 in 11 locations (Table 1). At harvest, fresh storage root yield and aboveground biomass (stem + leaves) were weighed separately. A net plot of 20 plants was harvested, excluding all plants on the border of a plot to avoid border effects. The storage root dry weight was calculated as a function of the fresh storage root yield and the dry matter content (Equation 3).

Dry matter content (DMC) in Ibadan was determined from 100 g of fresh roots of each plot grated and oven-dried at 65°C for 72 hours to constant weight. After the samples were cooled, their dry weights were recorded, and their respective DMC values were determined. For the other locations, the specific gravity method (Wholey & Booth 1979) was used to estimate the DMC per plot. The dry aboveground biomass was calculated as 1/3 of the fresh aboveground biomass one (Streck et al. 2014).

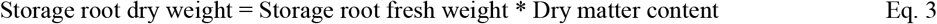

### Estimating genotype specific parameters

The DSSAT crop model contains genetic parameters at the levels of species, ecotype, and cultivar (Hoogenboom et al. 2019). For model evaluation, coefficients at the level of cultivars, referred as Genotype-Specific Parameters (GSPs), need to be calibrated. We also adjusted two ecotype coefficients directly involved in yield (Moreno-Cadena et al. 2020; Rankine et al. 2021).

The GSPs define differences among cultivars within a crop species (Hoogenboom et al. 2019). The 15 cultivar and 2 ecotype parameters incorporated within CSM-MANIHOT-Cassava that we calibrated for each cultivar are defined in Table 2. The prior distribution of each GSP was provided by Moreno-Cadena et al. (2020).

**Table 2.** Maximum (Max), minimum (Min), mean and coefficient of variation (CV) values of genotype-specific parameters calibrated for the 67 cassava cultivars in the MANIHOT-Cassava model.

| Coefficient | Fitted by | Max | Min | Mean | CV (%) |
| --- | --- | --- | --- | --- | --- |
| B01ND (°Cd) | Trial & Error | 1200 | 231 | 487 | 62 |
| B12ND (°Cd) | Trial & Error | 344 | 80 | 247 | 19 |
| B23ND (°Cd) | Trial & Error | 286 | 101 | 235 | 15 |
| B34ND (°Cd) | Trial & Error | 250 | 100 | 243 | 11 |
| BR1FX | Trial & Error | 3.8 | 2 | 2.5 | 13 |
| BR2FX | Trial & Error | 4 | 2 | 2.7 | 17 |
| BR3FX | Trial & Error | 3 | 1 | 2.1 | 17 |
| BR4FX | Trial & Error | 2 | 1.4 | 1.5 | 4 |
| LAXS (cm<sup>2</sup>) | Trial & Error | 920 | 401 | 754 | 19 |
| LLIFA (°Cd) | Trial & Error | 1800 | 600 | 1282 | 32 |
| SLAS (cm<sup>2</sup> g<sup>-1</sup>) | GLUE | 280 | 149 | 252 | 13 |
| LPEFR | GLUE | 0.3 | 0.2 | 0.3 | 17 |
| LNSLP | GLUE | 1.7 | 0.7 | 0.8 | 19 |
| NODWT (g) | GLUE | 6 | 1.7 | 3.3 | 32 |
| NODLT (cm) | GLUE | 4 | 1 | 2.9 | 27 |
| PARUE (g dry matter MJ<sup>-1</sup>) | Yield Matching | 2.6 | 1.4 | 1.8 | 21 |
| KCAN | Yield Matching | 0.8 | 0.5 | 0.6 | 15 |

Two methods were used to estimate GSPs. The first group of GSPs, including the phenological parameters and some growth parameters, were hand-calibrated following the trial and error procedure recommended by Hoogenboom et al. (2019) and used by Phoncharoen et al. (2021b). Simulations were repeated until we could not increase the agreement evaluated by the correlation (r) and the Root Mean Square Error values (RMSE) between the simulated and observed values. That method was adopted because the target variables were measured only in Ibadan 2021. The calibration procedure was as follows:

- Parameters defining plant phenology were calibrated in the respective order of their progress during growth. They included thermal time to first forking (B01ND), second forking (B12ND), third forking (B23ND), and fourth forking (B34ND). The target variables were the time of these events from planting (days after planting).
- The growth parameters measuring the number of branches per fork (BR1FX, BR2FX, BR3FX, and BR4FX) were directly measured. In case of a lack of data, the default value from the model GSPs file was applied.
- Leaf parameters, including the maximum leaf area (LAXS) and leaf life (LLIFA) were calibrated. The target variable was leaf area index (LAI) as computed in Eq. 2.

The second step of the calibration was done by the Generalized Likelihood Uncertainty Estimation (GLUE) method. The calibrated parameters were the specific leaf lamina area (SLAS), leaf petiole fraction (LPEFR), a slope for leaf production (LNSLP), node weight (NODWT) and internode length (NODLT). The GLUE method is one of the first methods to represent prediction uncertainty within the context of Monte Carlo analysis coupled with Bayesian estimation and propagation of uncertainty. The technique uses a likelihood function to measure the closeness-of-fit of modeled and observed data (He et al. 2010). The goal was to minimize the RMSE between simulated and observed values. The target variables were the aboveground biomass dry weight. The GLUE method was used because of its flexibility, ease of implementation as an R program, and its suitability for parallel implementation on distributed computer systems. It has previously been used to estimate GSPs for DSSAT crop models (Hoogenboom et al. 2019) such as wheat (Ibrahim et al. 2016; Li et al. 2018), maize (Mereu et al. 2019), and rice (Buddhaboon et al., 2018) as examples. The main steps of the GLUE program as described by Beven & Binley (1992), He et al. (2010), and updated by He et al. (2021), are summarized in Supplementary Methods.

After these two steps, if the storage root dry weight was systematically underestimated or overestimated across observations of a cultivar, the photosynthetically active radiation conversion factor (PARUE) and PAR extinction coefficient (KCAN) that are ecotype parameters were adjusted manually to remove the bias. Then, the GLUE method was run again as defined above to estimate the GSPs using new values of PARUE and KCAN.

## Statistical analysis

### Model performance analysis

The performance of the calibrated CSM-MANIHOT-Cassava model was analyzed using the correlation (r), the Root Mean Square Error (RMSE, Wallach & Goffinet 1987), the normalized root mean square error (nRMSE, Loague & Green 1991) and the index of agreement (d, Willmott 1982). These four statistical parameters were analyzed separately for storage root dry weight. The parameters r (Equation 4), RMSE (Equation 5) and nRMSE (Equation 6) measure the deviation of the simulated from the observed values. The nRMSE is the RMSE value divided by the observed mean. The simulation is considered excellent with nRMSE < 10%, good in the 10-20% range, fair in the 20-30% range, and poor if nRMSE is > 30% (Harb et al. 2016; Rankine et al. 2021). The index of agreement, *d*, (Equation 7) was added because it incorporates both bias and variability. The *d* value varies between 0 and 1. A value of 1 indicates a perfect match, and 0 indicates no agreement at all.

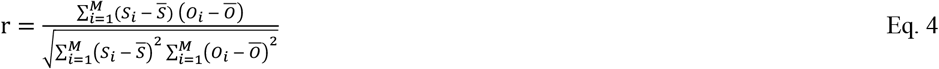

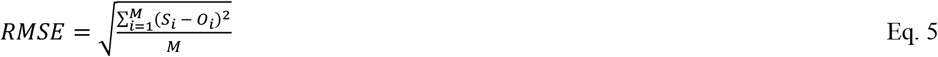

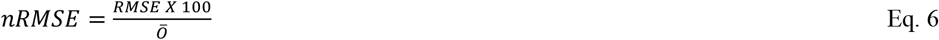

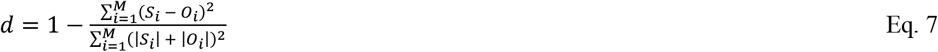

*M =* number of observations

*S*_*i*_ = simulated value

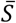 = mean of the simulated values

*O*_*i*_ = observed value

Ō = mean of the observed values

### Model sensitivity analysis

The main objective was to identify variables explaining deviations between model predictions and experimental data. The hypothesis was that the differences between the simulated and the observed values could be due either to a bad calibration of the GSPs (leading to a genotype effect on model deviation) or to the fact that some parameters of the environment were not well accounted for in the model (leading to an environment effect on model deviation) or to an interaction between these two causes (leading to a genotype by environment interaction effect on model deviation).

We analyzed deviations using the Additive Main effect and Multiplicative interaction (AMMI) model implemented in the statgenGxE R package (van Rossum et al. 2022). The AMMI model separates additive main effects and multiplicative interaction effects. The model analyses the additive effects with classical analysis of variance (ANOVA) then applies principal component analysis (PCA) to the residual interaction portion of variation (Hongyu et al. 2014; Nowosad et al. 2016; Yan & Rajcan 2002).

The AMMI model is:

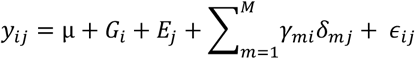

where:

- *M* was the number of principal components included,
- *y*_*ij*_ was the deviation between observed versus simulated dry storage root yield for genotype *i* in environment *j*. That is, *y*_*ij*_ = Observed storage root dry weight – simulated storage root dry weight,
- *μ* was the general mean,
- *G*_*i*_ was the genotypic effect,
- *E*_*j*_ was the environmental effect,
- *γ*_*mi*_ was the genotypic score for principal component *m* and genotype *i*,
- *δ*_*mj*_ was the environment loading for principal component *m* and environment *j*,
- *ϵ*_*ij*_ was the residual.

For example, if after calibration, observed yields of a genotype across environments are consistently higher than those simulated by the CSM-MANIHOT-Cassava model, *G*_*i*_ will be positive. Likewise, if observed yields across genotypes in an environment are consistently higher than those simulated by CSM-MANIHOT-Cassava, *E*_*j*_ will be positive. Conditions for *γ*_*mi*_ and *δ*_*mj*_ to be non-zero would require that certain aspects of the genotype interact with certain aspects of the environment to cause a bias in CSM-MANIHOT-Cassava simulations. Explanatory variables considered for these analyses included weather data, soil data, and additional phenotypic data, encompassing diseases such as Cassava Bacterial Blight incidence and severity (CB_), Cassava Green Mite severity (CGM), Cassava Mosaic Disease incidence and severity (CMO), as well as leaf area (LeafArea) and plant morphology, such as Plant forking number (branchP), Stem number (stem_nu), Stem diameter (stem_D), and Plant height (plant_H). The methods used for obtaining additional data were consistent with those described in Rabbi et al. (2014), Rabbi et al. (2017), and Wolfe et al. (2016).

## Results

### Variation among optimized GSPs

The cultivars presented variation in the optimized parameters controlling phenological events and growth (Table 3). For the phenology, the first forking (B01ND) time presented the highest coefficient of variation (CV = 62%). The second, the third, and the fourth forking time showed substantially lower CVs. The number of branches for the first forking was between 2 and 4, and some cultivars had 4 branches in the second branching.

**Table 3.**
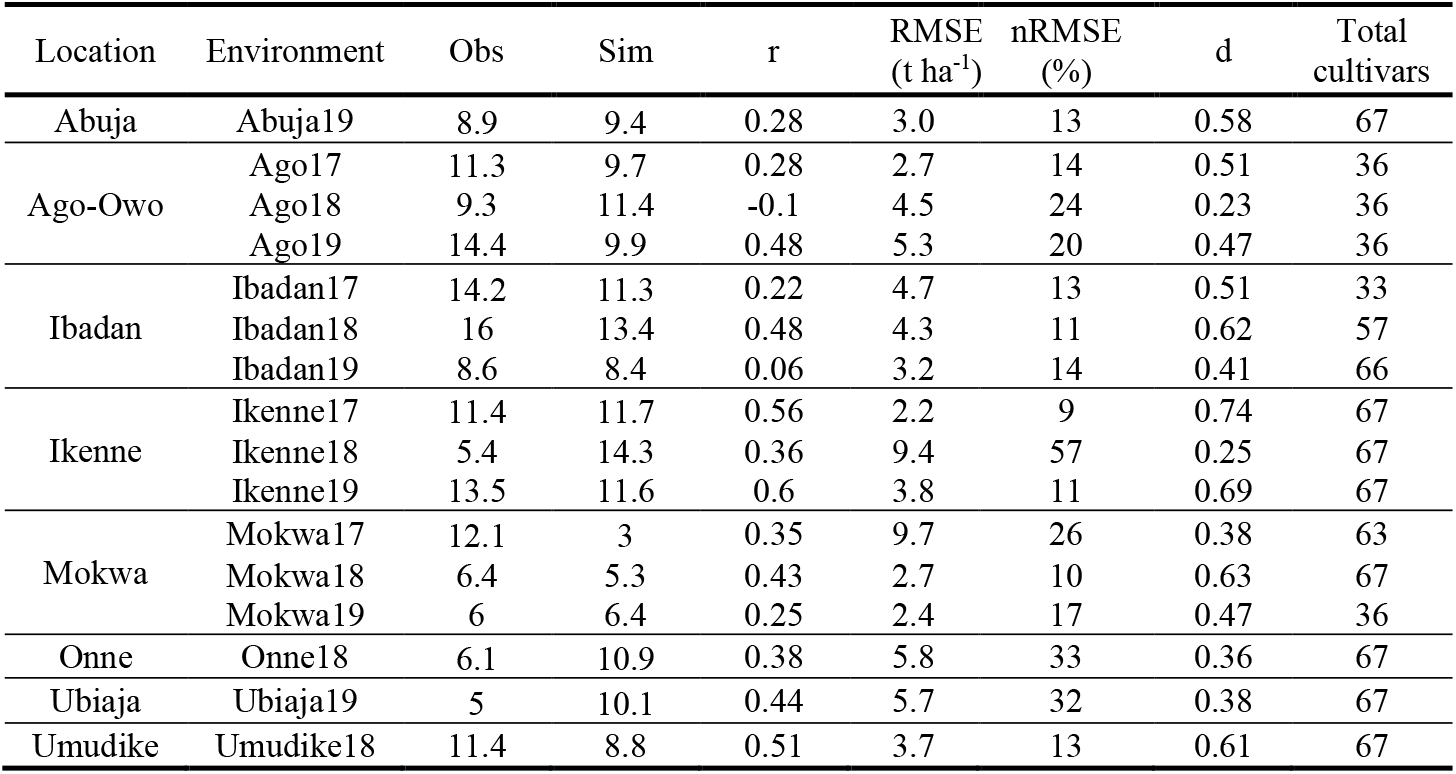
Agreement between simulated (Sim) and observed (Obs) mean storage root dry weight of the cultivars across environments for MANIHOT-Cassava calibration.

Parameter variation was observed at both the leaf and internode. The leaf production and functioning parameters, such as LAXS and SLAS presented CV values of 19% and 13%, respectively. Similarly, the internode weight (NODWT) and length (NODLT) showed CVs of 32% and 27%, respectively, which were the highest CVs among the GLUE-optimized parameters. Finally, among the parameters affecting photosynthetic activity, PARUE and KCAN had 21% and 15% CV values, respectively

### Model performance

#### Performance by cultivar

The analysis included the 67 cultivars evaluated across 16 environments. For each cultivar we calculated the Pearson correlation r of model prediction and observation across the environments, as well as the RMSE and d statistic. The GLUE estimated GSPs improved the accuracy of the dry storage root yield simulated by the model (Figure 3). The correlation coefficient increased from – 0.03 to +0.08, the RMSE dropped from 21 t ha^-1^ to 5 t ha^-1^ while d increased from 0.23 to 0.44. All these changes were highly significant using a paired t-test.

**Figure 3.**
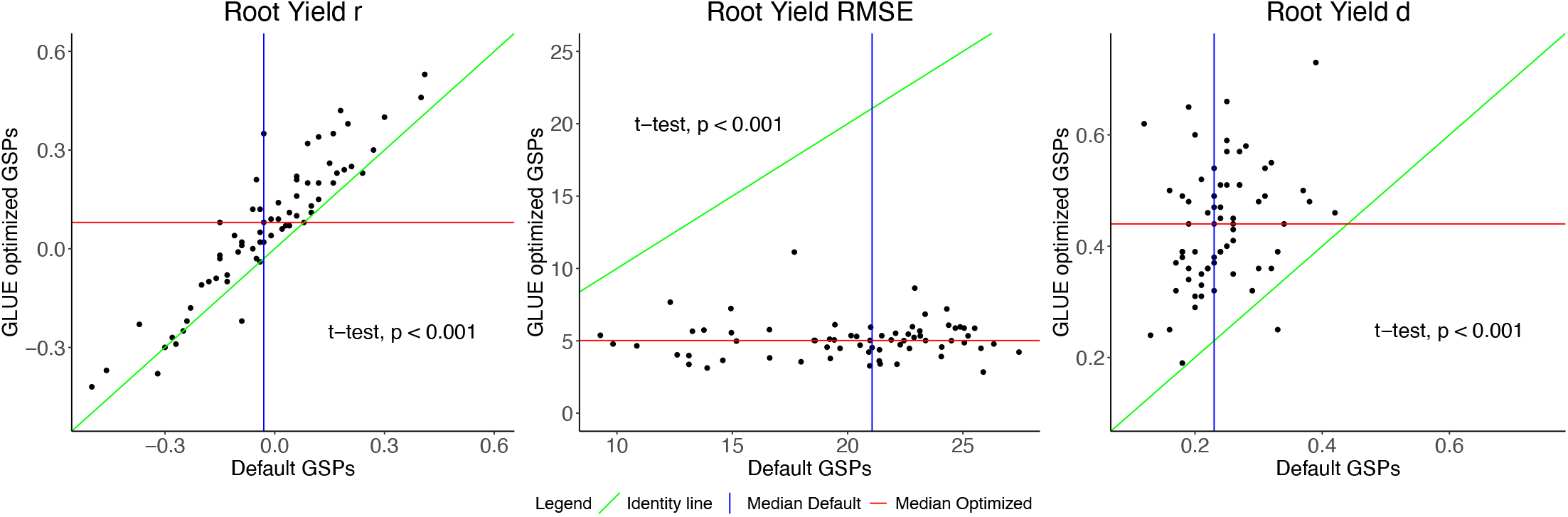
Comparison of the Pearson correlation coefficient (r), the root mean squared error (RMSE), and the index of agreement (d) values for the MANIHOT-Cassava predicted storage root dry weight before and after using General Likelihood Uncertainty Estimation (GLUE) method for Genotype-Specific Parameters (GSPs) estimation. Each point represents one cultivar.

**Figure 4.**
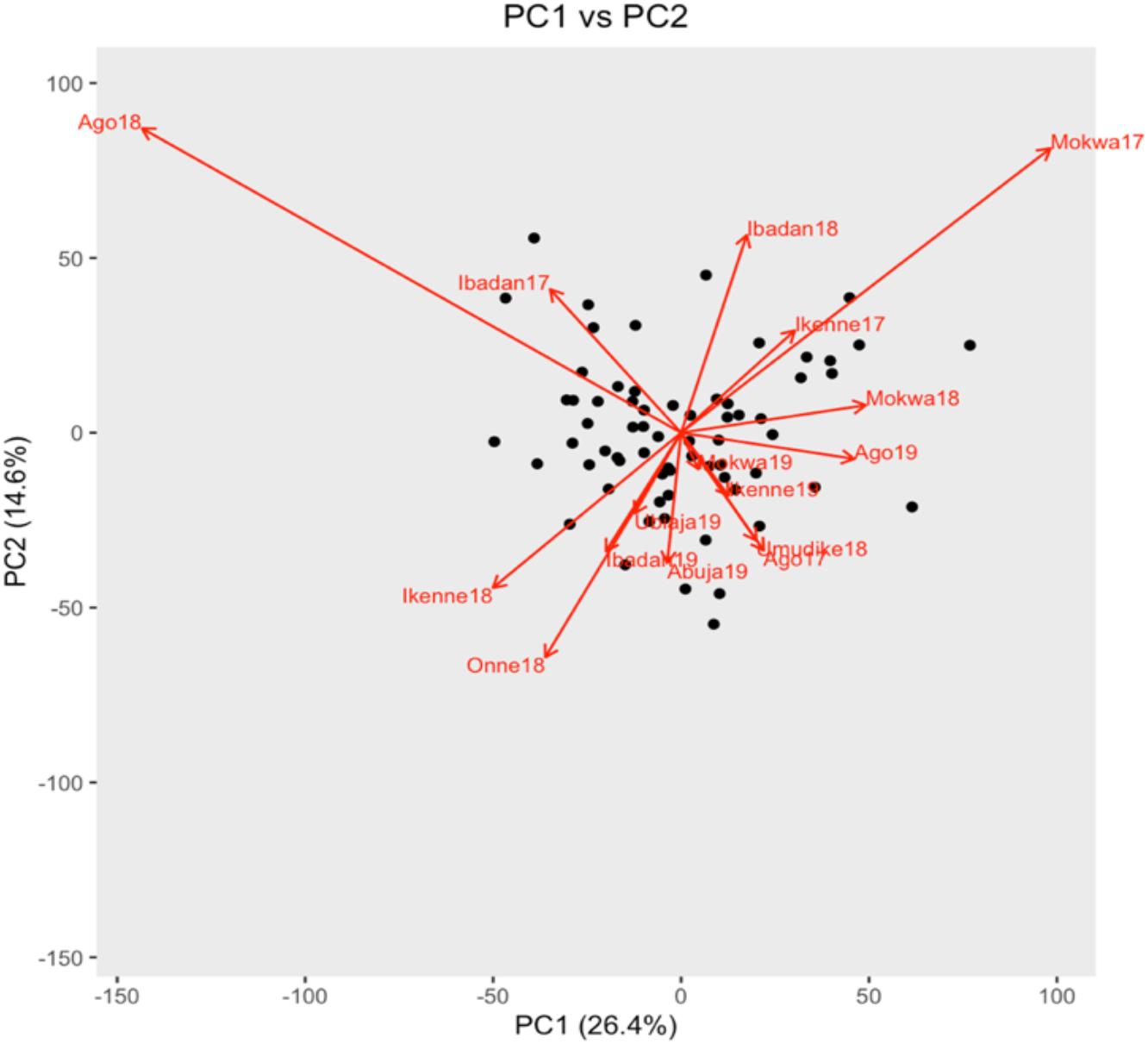
Biplot of genotypes and environment interaction with PC1 and PC2. Genotypes are black points, and environments are red arrows.

#### Performance by environment

The r, RMSE, and d statistics could also be calculated across cultivars within environments. In this per-environment analysis, we also observed reasonable agreement between the simulated and the observed storage root dry weight. A total of 69% of the environments had an nRMSE ≤ 20% (considered excellent to good). Some environments such as Ikenne17 and Mokwa18 were very well simulated with nRMSE values of 9% and 10%, RMSE of 2.2 and 2.7 tha^-1^ and d values of 0.74 and 0.63 respectively (Table 4). Equally, substantial deviation of the model was observed in some environments such as Ikenne18 and Onne18 (nRMSE ≥ 30%). Given this observation, statistical analyses of simulation results were performed to determine the causes of the model deviations.

**Table 4.**
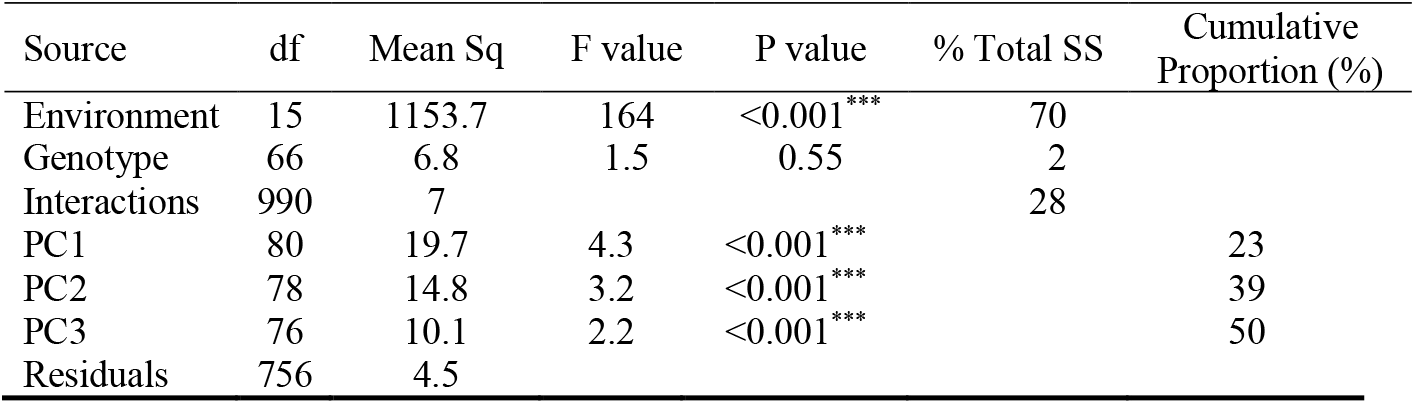
Analysis of variance for the model storage root dry weight (t ha^-1^) deviation (observed – simulated) using the AMMI model. With degrees of freedom (df), mean square (Mean Sq), the F value and P value and the proportion of sums of squares (% Total SS).

| Source | df | Mean Sq | F value | P value | % Total SS | Cumulative Proportion (%) |
| --- | --- | --- | --- | --- | --- | --- |
| Environment | 15 | 1153.7 | 164 | <0.001*** | 70 |  |
| Genotype | 66 | 6.8 | 1.5 | 0.55 | 2 |  |
| Interactions | 990 | 7 |  |  | 28 |  |
| PC1 | 80 | 19.7 | 4.3 | <0.001*** |  | 23 |
| PC2 | 78 | 14.8 | 3.2 | <0.001*** |  | 39 |
| PC3 | 76 | 10.1 | 2.2 | <0.001*** |  | 50 |
| Residuals | 756 | 4.5 |  |  |  |  |

### Genotype and environment effects on model prediction error

The objective of the AMMI analysis was to identify the possible causes that could be ascribed to cultivars or environments of poor model performance in predicting root yield. The analysis was applied to the difference between observed and simulated storage root dry weight (observed – simulated = “O-S”). The AMMI model evaluated 66 genotypes across 16 environments. One genotype out of the 67 was not included because it had more than 10% missing data. The ANOVA analysis revealed a significant effect only for the environment explaining model error (Table 5). The effect of the genotype and environment interaction was also highly significant (p < 0.001). The first three interaction principal components (PCs) explained 50% of the GEI variation.

The biplot of model prediction errors indicated that the environment effects were strongest at Ago18 and Mokwa17 whose effects were opposite on the component of GEI explained by PC1 (Figure 5).

**Figure 5.**
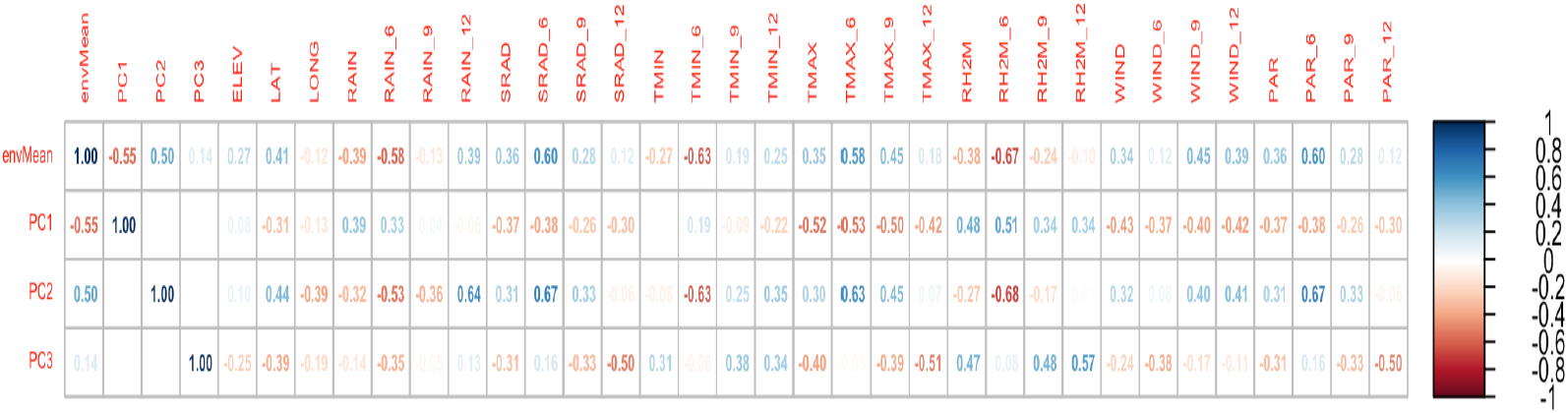
Correlation between the mean value of the model deviation (“Observed – Simulated”) by environment (envMean), the environment loading values on PC1, PC2 and PC3, and weather parameters The suffix number (“_x”) defines the months after planting, e.g., TMAX_6 is the maximum temperature at 6 MAP. Abbreviations are: ELEV: location altitude (m), LAT: location latitude, LONG: location longitude, RAIN: rainfall from planning to harvest; SRAD: solar radiation, TMIN: minimum daily temperature, TMAX: daily maximum temperature, RH2M: relative humidity, WIND: average wind speed at 2m, PAR: Photosynthetically active radiation.

### Effects of environment characteristics on model prediction error

To identify characteristics of environments that affected model accuracy, we correlated weather and soil profile data with the AMMI analysis output including the mean value of model deviations (i.e., “O-S”) by environment (envMean) and the environment loading values on the three PCs.

Many input weather parameters affected the model performance (Figure 6). Indeed, the solar radiation, maximum temperature, wind, and daily photosynthetic radiation were positively correlated with the deviations while rainfall and relative humidity were negatively correlated (p-values < 0.01). Root yields in dry (low rain) environments marked by high temperatures and high solar radiation were underestimated by the model, while yields in wet (high rain) environments with low temperatures and low solar radiation were overestimated (Figure 7). These weather parameters affected estimation more in the first six months of plant growth: parameters at 6 months after planting (parameter abbreviation with “_6”) had the strongest correlations.

**Figure 6.**
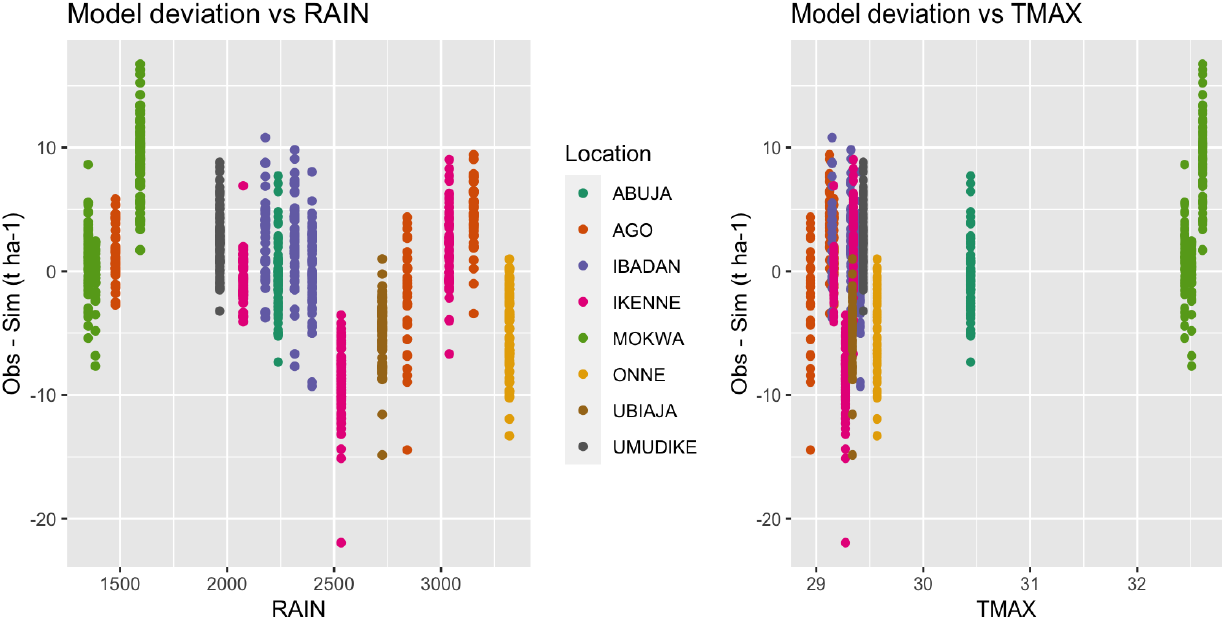
Scatterplot of the mean model deviation (“Observed – Simulated”) against the annual total rain by environment (RAIN) and the annual mean maximum temperature by environment (TMAX)

**Figure 7.**
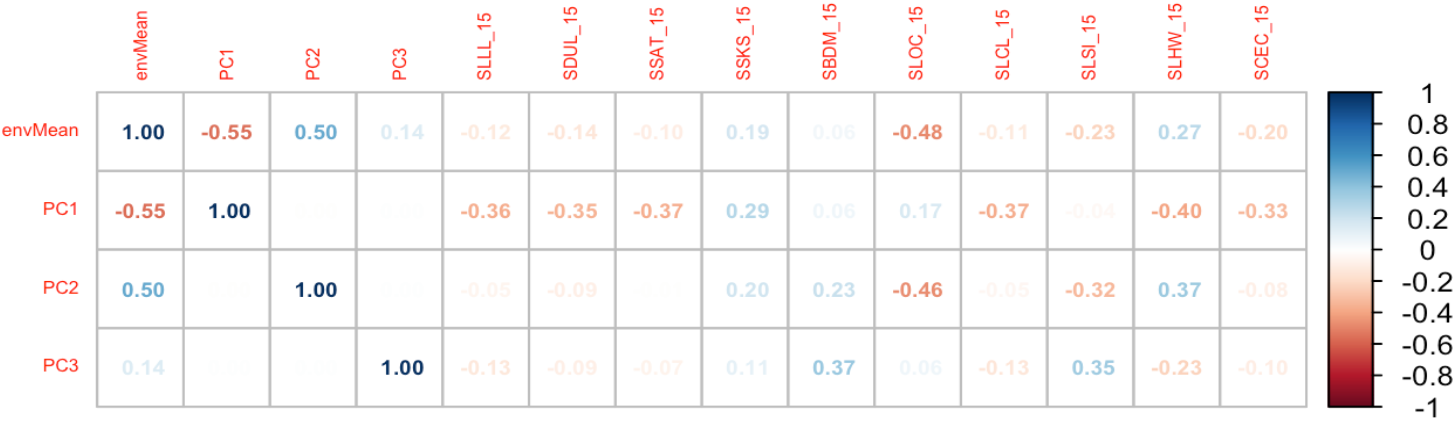
Correlation between the mean value of the model deviation (“Observed – Simulated”) by environment (envMean), the environment loading values on PC1, PC2 and PC3, and soil profile parameters. Soil parameters abbreviations are SLLL: Lower limit of plant extractable soil water (cm^3^), SDUL: Drained upper limit (cm3), SSAT: Upper limit, saturated (cm^3^), SSKS: Saturated hydraulic conductivity (cm/h), SBDM: Bulk density (g/cm^3^), SLOC: Organic carbon (%), SLCL: Clay (%), SLSI: Silt (%), SLHW: pH in water, SCEC: Cation exchange capacity (cmol/kg).

Subsoil characteristics at 15 and 100 cm were strongly correlated, and we only present results at 15cm. Only the soil organic carbon (SLOC), significatively affected MANIHOT-Cassava performance showing a negative correlation with the model deviations (Figure 8).

## Discussion

### Variation in cultivar GSPs

Crop Growth Models are usually calibrated in non-stress environments. These environments enable the expression of the cultivar’s inherent growth and potential without confounding them with stress factors. Our objective here, however, was to calibrate with breeding trial data to avoid requiring the breeding program to allocate resources to model calibration experiments. We assume that the lack of ideal environments can be compensated for by having data from many environments.

Across the 15 GSPs, we used three calibration approaches: Trial & Error, GLUE, and Yield Matching (Table 3). For the Trial & Error calibration we only used data from the Ibadan 2021 trial for which we had estimates of leaf area and of branch phenology and number of forks. These parameters were fixed in part because we found that the GLUE approach could not handle more than the five GSPs that were calibrated with it. Because GLUE samples from the prior distribution of parameters it is not super-efficient at narrowing in on values with maximum posterior density (He et al. 2010). Recent efforts have been made to parallelize GLUE (Berton Ferreira 2024) to make it more computationally feasible. Finally, we set PARUE and KCAN to come close to matching the average yield of the cultivars. This approach was chosen to reduce the impact of genotype main effects on model calibration and instead capture sources of genotype by environment interaction. The approach is not unlike post hoc adjustments for harvest index to match harvested yield to model biomass simulation (as might have been done, for example, in Wellens et al. 2022). One consequence of that choice is that model deviations were barely affected by genotype (Table 5) so that we were not able to explore effectively genotype characteristics that might affect model performance. Adjusting to match genotype main effect, however, arithmetically could not affect the correlation for a genotype between simulation and observation across environments. Thus, the model improvement in that correlation from a median across cultivars of -0.03 for the default parameters to 0.08 was due to calibration of the other parameters. While the optimized correlation is not high, the improvement was statistically significant (Figure 3).

Across calibration approaches, we found evidence of variability between the IITA cultivars. Concerning plant phenology, the CV among cultivars was 62% for thermal time to first branching. This parameter establishes the difference between early- and late-branching genotypes (Moreno-Cadena et al. 2020). Similar results were obtained on the elite cassava cultivars from the National Crops Resource Research Institute breeding program of Uganda (Ibrahim et al. 2020) using standard statistical analysis. For Thai genotypes, Phoncharoen et al. (2021b) obtained a very high variation 97%, unlike the Jamaica genotypes where Rankine et al. (2021) found a CV of only 10%. Branching is a significant event in the cassava’s life cycle. Phenology not only determines the rate of production of leaves and the architecture of the plant but also accounts for the plant’s ability to flower, because an inflorescence is always coupled with branching (Halsey et al. 2008; Jennings & Iglesias 2001). Thus, the level and timing of branching influence flower availability and fertility (Ibrahim et al. 2020). The B12ND, B23ND and B34ND parameters and their corresponding number of branches (BRnFX) could be interpreted because not all the cultivars showed a fork when the data was collected between 5 and 6 months after planting. For those cultivars, the model default values were used (Moreno-Cadena et al. 2020).

There was also variation among the cultivars for the leaf parameters. These results indicate that the germplasm of IITA was characterized by high variability of leaf production, growth, as well as leaf functioning as shown in the PARUE and KCAN parameters. The importance of getting leaf parameters well calibrated is that the MANIHOT-Cassava model uses a surplus of assimilate approach to determine when to allocate to storage roots. We were only able to measure LAI in one environment. More data on cassava LAI across studies might improve model performance. At least two different data collection times during the growing season are needed to calibrate the genetic parameters (Banterng et al. 2009; Phoncharoen al. 2021b; Rankine et al. 2021; Suriharn et al. 2007).

The individual node weight (NODWT) showed a high variation across the target germplasm followed by node length (NODLT) and leaf production slope (LNSLP). According to Moreno-Cadena et al. (2020), NODWT and LNSLP are both essential GSPs affecting plant total biomass with the latter being particularly important in warm environments. The NODWT parameter is important because it affects aboveground photosynthate demand which must be satisfied before the remaining assimilates are allocated to the storage roots. A high value for NODWT can lead to a decrease in storage root dry weight (Moreno-Cadena et al. 2021; Phoncharoen et al. 2021b).

### Comparison to other model calibrations

Model calibration across 67 cultivars with final fine calibration using the GLUE method improved the prediction of the storage root dry weight. Strong model improvements for the RMSE and d statistics were no doubt partially due to the yield matching approach we used to calibrate PARUE and KCAN. The improvement in r that we observed was due to other GSP adjustments, however. Similar methodology was successfully used to calibrate the CSM-CERES-Wheat and CSM-CERES-Maize models for winter wheat and maize grain yield simulation, achieving RMSE of 0.27 and 1.57 t ha^-1^, respectively, and d values of 0.98 and 0.85, respectively (Ibrahim et al. 2016; Mereu et al. 2019). However, RMSE values of 2.19 and 3.72 t ha^-1^ were obtained for durum wheat and common wheat grain yield simulation, respectively, which are similar to the results we obtained in some environments (Mereu et al. 2019). The cassava storage root yield for six locations in Nigeria was simulated with an RMSE of 4.9 t dry matter ha^−1^ using the LINTUL-Cassava model (Adiele et al. 2021). The nRMSE values obtained by environment were similar to those reported by Phoncharoen et al. in two diferrent studies (2021a; 2021b). Finally, Wellens et al. (2022) achieved similar normalized RMSE in the range of 11% to 33%.

Finally, our model fit statistics could have been affected by the fact that we did not have explicitly measured local soil and weather data. Lack of accurate local soil profile data and specific topsoil chemical analysis data could lead to the model prediction errors. To assess this possibility further, we calibrated the model under both nitrogen-limited and non-nitrogen-limited conditions; phosphorus not being an important limiting factor in the CSM-MANIHOT-Cassava model (Moreno-Cadena et al. 2021). Model accuracy was better under non-nitrogen-limited conditions. The weather data we used came from NASA sources; a local weather station would have been preferred. In future research we will also seek to use CHIRPS rainfall data (Funk et al. 2015).

### Model variables to investigate further

To our knowledge no cassava study has separately calibrated a model for as many cultivars as we have here. We believe that the extensive set of cultivars we calibrated lends robustness to our conclusions on model performance. The analysis of model prediction errors showed that root yield prediction error was linked to the input weather parameters SRAD, TMAX, wind, rain, and RH2M (Figure 7). The fact that genotypes clustered together when plotted by environments also reflects the importance of environment on model prediction error (Figure 5). The correlation between weather parameters and model prediction error could be used to classify the environments and identify physiological relationships to be improved in the model.

The underestimated environments were characterized by low rain resulting in low relative humidity. The model integrates rainfall as plant extractable soil water by subtracting surface runoff, drainage, and soil evaporation (Hoogenboom et al. 2019). As root water uptake decreases, stomatal conductance decreases through stomatal closure, which reduces carbon assimilation and transpiration as well as nutrient absorption (Moreno-Cadena et al. 2020). In addition, prior studies with the model showed that under warm temperatures water availability could be a limiting factor for cassava root yield (Moreno-Cadena et al. 2021; Phoncharoen et al. 2021a; Rankine et al. 2021). Cassava stomata are sensitive to vapor pressure deficit (VPD) and reduce gas exchange more than other crops, which can limit photosynthesis while also saving water (Oguntunde & Alatise 2007; Vongcharoen et al. 2018). The model currently incorporates a drought stress factor that integrates a VPD factor for photosynthesis. Our calibration study suggests that this stress factor might need to be adjusted. The improved cultivars from the IITA breeding program are selected for high performance including in water stress environments in the north of Nigeria. These cultivars could therefore be more tolerant than assumed by the model. A drought condition assumed by the model could still allow normal root growth for these improved genotypes. At the same time, increased rainfall led to higher predicted storage root dry weight but did not affect the observed values. The model requires defining thresholds for water and nitrogen stress, leaf and root senescence, and root water uptake (Moreno-Cadena 2021). These are currently species coefficients that were not modified by our study. Modifying these settings could improve model performance in more extreme wet or dry environments.

Finally, the IITA improved cultivars might not follow the photosynthetic assimilate allocation strategy of CSM-MANIHOT-Cassava model which feeds the storage root growth with the remaining assimilate after having satisfied the aboveground sink. The spill over model is elegant and has scientific support (Cock & Connor 2021) but is not a consensus (e.g., Gray 2000). Several studies showed that improved cassava cultivars preferentially allocate photosynthetic assimilates to storage roots after their initiation (Lahai et al. 1999; Turyagyenda et al. 2013). The partitioning of the assimilate and the dynamics of the source-sink carbon allocation during the life cycle of the plant are important phenomena to understand to better simulate cassava’s growth. In a study contrasting two cassava varieties, one genotype favored stem growth while the other favored the storage roots once they were initiated (Chiewchankaset et al. 2022). Finally, a given cultivar need not be limited to a single allocation strategy but that, at a stage of its growth and according to its genotype and environmental factors, the cultivar adopts different mixtures of strategies.

## Data Availability

The datasets and scripts used in the analyses presented here can be found on GitHub: https://github.com/Opamelas83/RDSSATMICH.

## Author Contributions

PO, J-LJ conceived and designed the study; PO, IR, SK Implemented field trials and methodology, PO, J-LJ and PM-C curated the data, performed data analyses, and wrote the manuscript, PO, J-LJ, PM-C and GH edited the manuscript; J-LJ and SK Provided overall coordination and leadership.

## Acknowledgments

The authors thank the Bill & Melinda Gates Foundation for their financial support. Thanks to the staff of the International Institute of Tropical Agriculture (IITA) of the cassava breeding program that assisted for implementing the field experiments and collecting the data.

## Supplementary Data

**Supplementary Figure 1:**
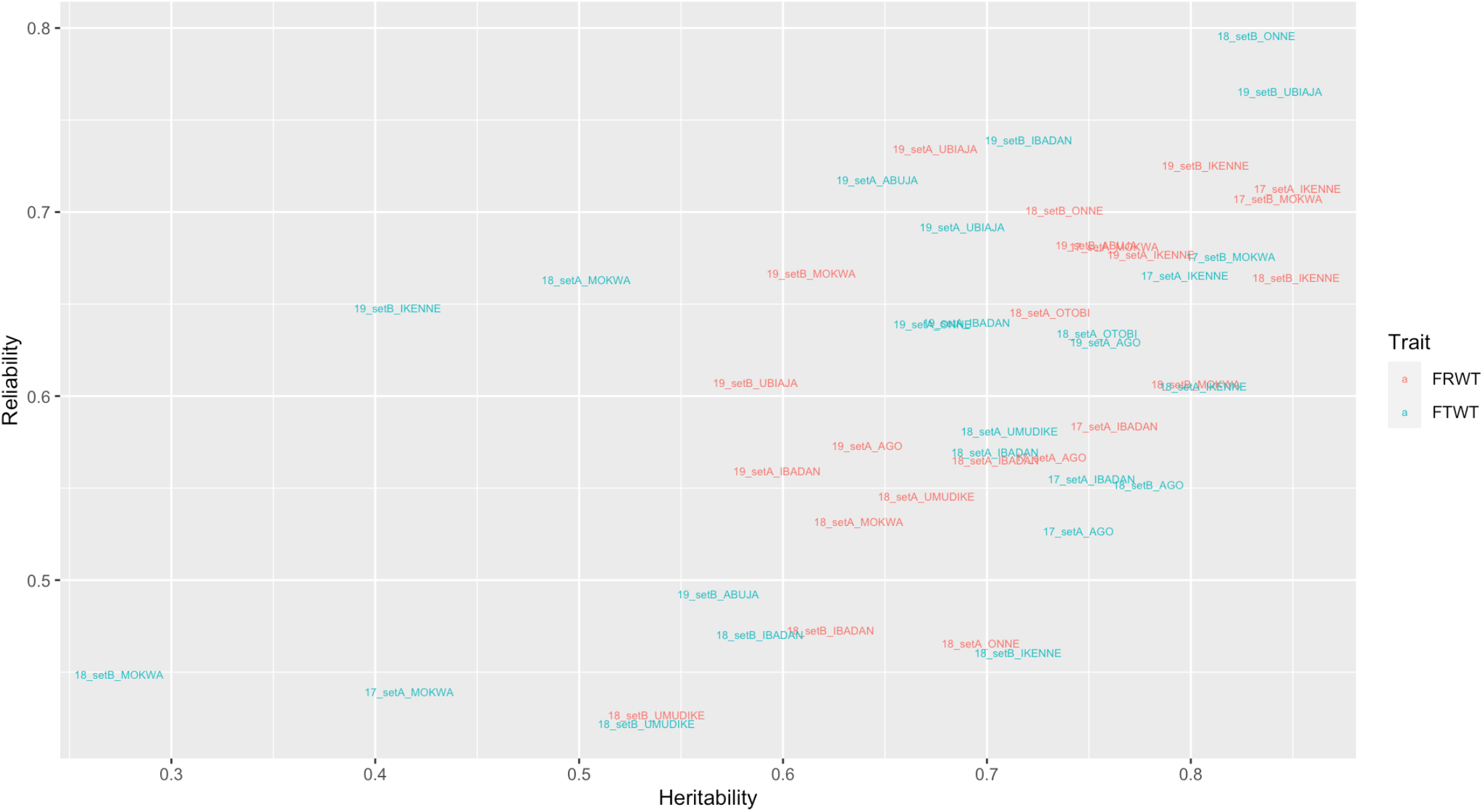
Plot of the reliability and heritability of the single trials used for the study. FRWT= Fresh storage root weight; FTWT= Fresh above ground biomass weight

**Supplementary Table 1:**
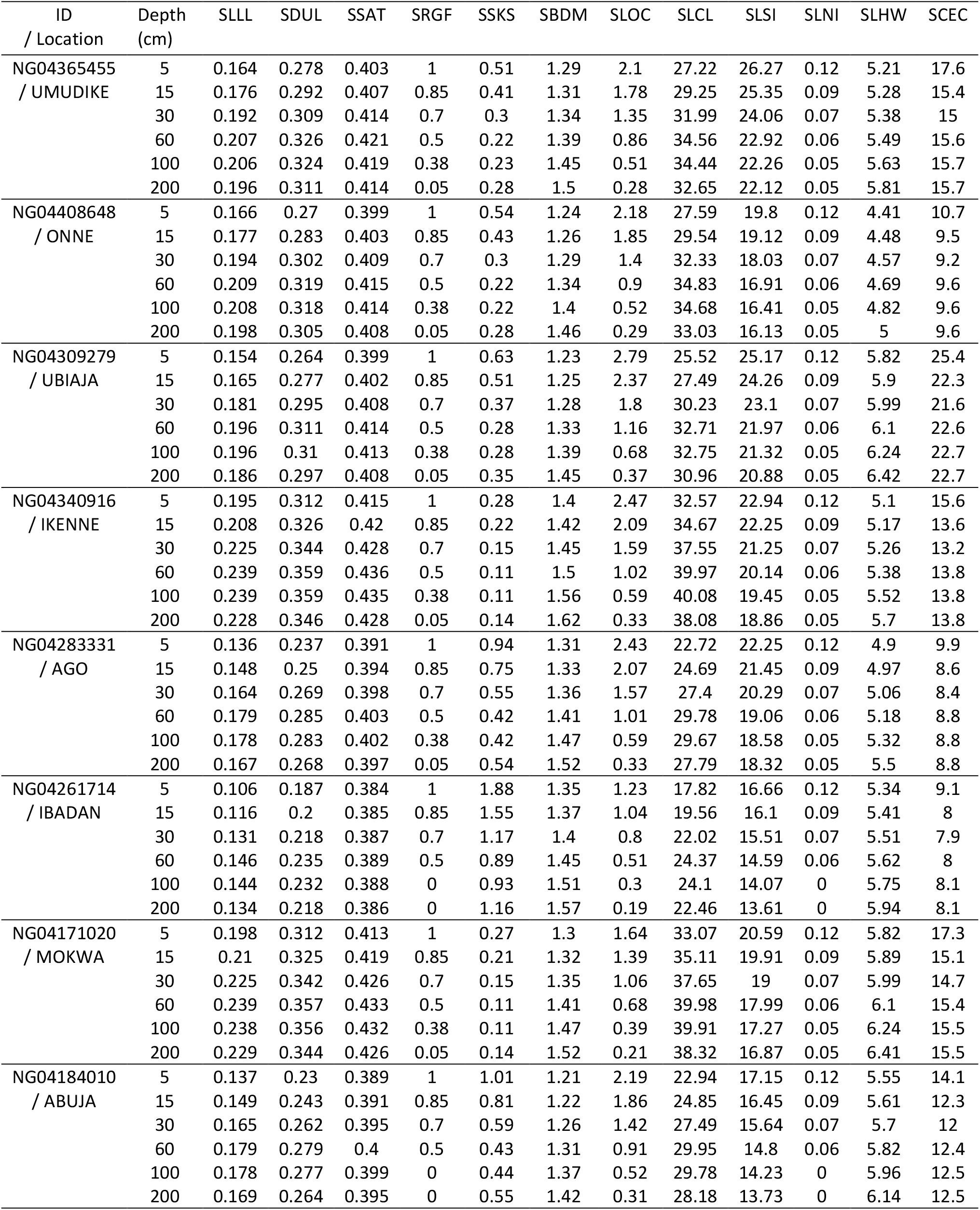
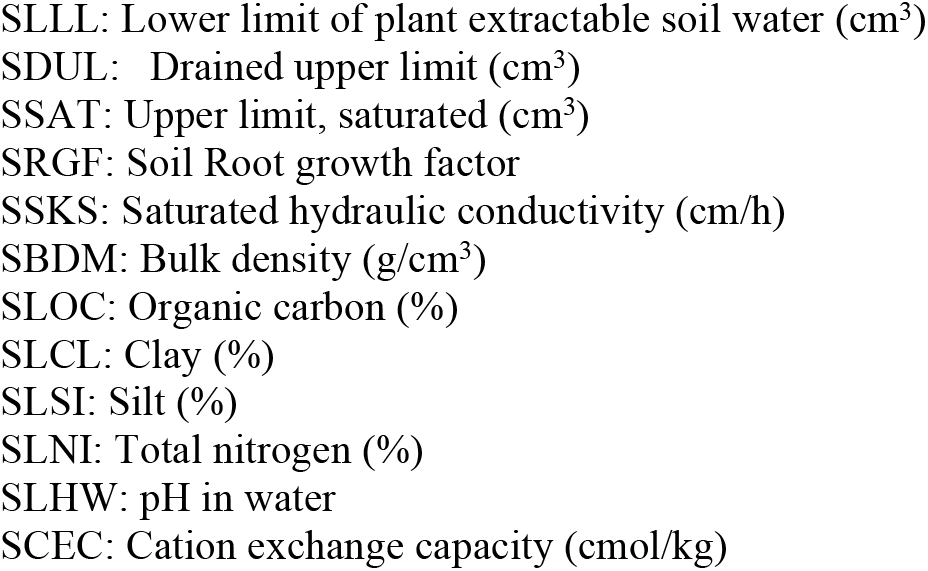
Physical and chemical soil properties in 8 locations.

## Supplementary Methods

The Generalized Likelihood Uncertainty Estimation (GLUE) program as described by Beven & Binley (1992), He et al. (2010), and updated by He et al. (2021).

### a. Develop prior parameter distributions

Random sampling, based on the prior distribution of parameters, is generally implemented in GLUE. The ranges of the parameters were defined and stored in a file formatted for the GLUE program under DSSAT 4.8.0.

### b. Generate random parameter sets from the prior parameter distributions

The parameters sets were generated using a Monte Carlo sampling of distributions implemented in a MATLAB program. From the point of view of Monte Carlo sampling in the GLUE method, more parameter sets lead to more stable results (Li et al., 2018). Thus, a total of 10,000 (the minimum recommended is 3,000) random parameter sets were generated from the prior distribution.

### c. Run the MANIHOT-Cassava model

The MANIHOT-Cassava model was run with each parameter set generated above by introducing the set into the standard genetic input file (cultivars file). For each run, the program identified all experiments in the DSSAT cassava data files involving the accession selected and these were used in the coefficient estimation process. The crop growth simulation outputs of dry aboveground biomass for each parameter set were tabulated for use in the GLUE likelihood calculations.

### d. Calculate the likelihood values and probability

The observed data on dry aboveground biomass were used along with the corresponding simulated outputs to compute the likelihood value. The Likelihood Function is the product of these individual likelihood values. The Gaussian likelihood function used in this study was recommended by He *et al*. (2010):

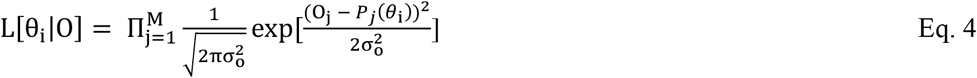

*L*[*θ*_*i*_|*O*] = likelihood value of parameter set i, given the set of observations

*θ*_*i*_ = i^th^ parameter set (I = 1, 2, 3, …N). N = 10,000 in our case.

*O*_*j*_ = j^th^ observation

*P*_*j*_(*θ*_*i*_) = model output for the j^th^ environment under parameter set *θ*_i_

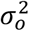= variance of model error M = number of observations.

The posterior probability *p*(θ_*i*_), of each parameter set θ_*i*_ is computed with the Bayesian equation as follows:

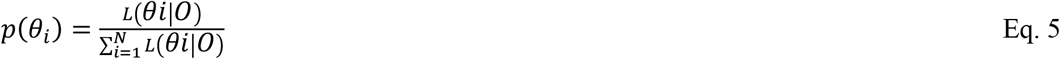

### e. Construct posterior distribution and statistics

The pair of parameter sets and probabilities [θ_*i*_, *p*(θ_*i*_)] were used to compute the mean, and the variance of each parameter Σ_i_:

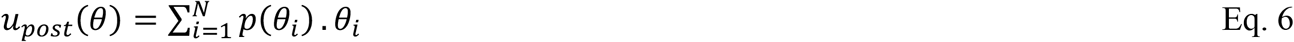

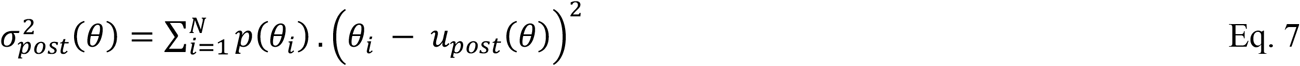

*u*_*post*_(θ_*i*_) = mean of the posterior distribution

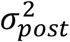*=* variance of the posterior distribution

