## Supplemental Data and Methods for "Evaluating a Cassava Crop Growth Model by Optimizing Genotypic-Specific Parameters Using Multi-environment Trial Breeding Data"

Supplementary Data

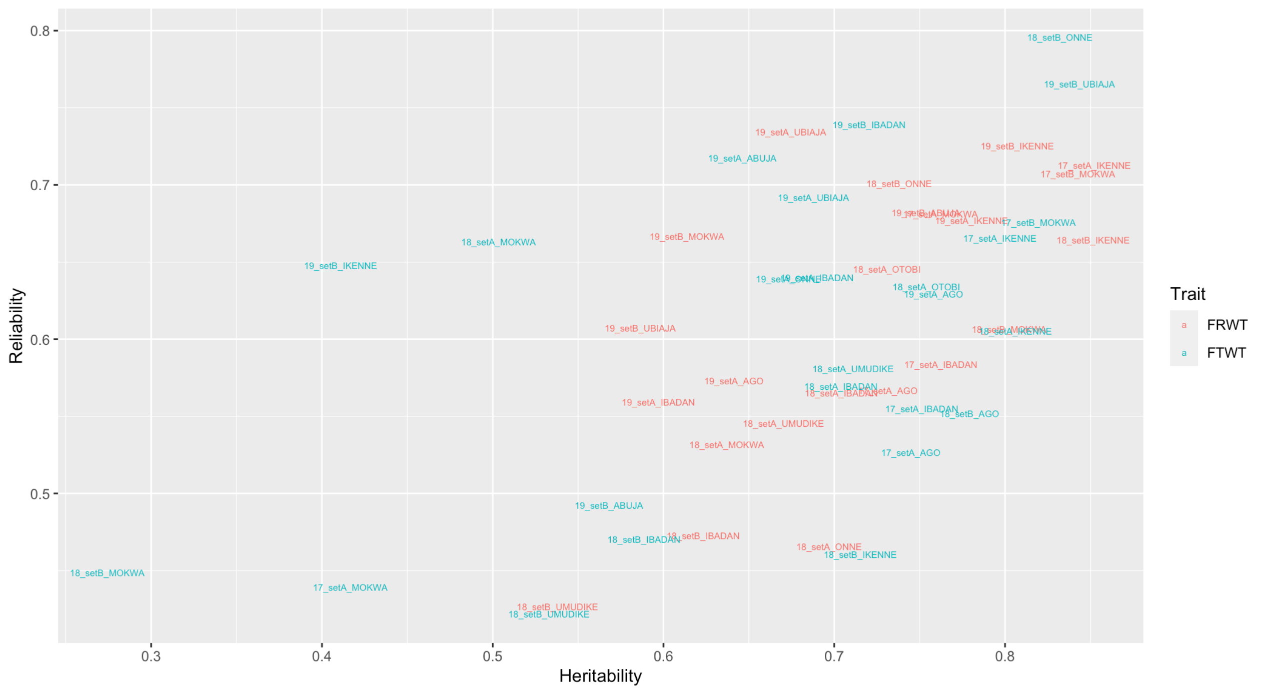


Supplementary Figure 1: Plot of the reliability and heritability of the single trials used for the study.

FRWT= Fresh storage root weight; FTWT= Fresh above ground biomass weight

Supplementary Table 1: Physical and chemical soil properties in 8 locations

| ID / Location | Depth (cm) | SLLL | SDUL | SSAT | SRGF | SSKS | SBDM | SLOC | SLCL | SLSI | SLNI | SLHW | SCEC |
|---|---|---|---|---|---|---|---|---|---|---|---|---|---|
| NG04365455 / UMUDIKE | 5 | 0.164 | 0.278 | 0.403 | 1 | 0.51 | 1.29 | 2.1 | 27.22 | 26.27 | 0.12 | 5.21 | 17.6 |
| | 15 | 0.176 | 0.292 | 0.407 | 0.85 | 0.41 | 1.31 | 1.78 | 29.25 | 25.35 | 0.09 | 5.28 | 15.4 |
| | 30 | 0.192 | 0.309 | 0.414 | 0.7 | 0.3 | 1.34 | 1.35 | 31.99 | 24.06 | 0.07 | 5.38 | 15 |
| | 60 | 0.207 | 0.326 | 0.421 | 0.5 | 0.22 | 1.39 | 0.86 | 34.56 | 22.92 | 0.06 | 5.49 | 15.6 |
| | 100 | 0.206 | 0.324 | 0.419 | 0.38 | 0.23 | 1.45 | 0.51 | 34.44 | 22.26 | 0.05 | 5.63 | 15.7 |
| | 200 | 0.196 | 0.311 | 0.414 | 0.05 | 0.28 | 1.5 | 0.28 | 32.65 | 22.12 | 0.05 | 5.81 | 15.7 |
| NG04408648 / ONNE | 5 | 0.166 | 0.27 | 0.399 | 1 | 0.54 | 1.24 | 2.18 | 27.59 | 19.8 | 0.12 | 4.41 | 10.7 |
| | 15 | 0.177 | 0.283 | 0.403 | 0.85 | 0.43 | 1.26 | 1.85 | 29.54 | 19.12 | 0.09 | 4.48 | 9.5 |
| | 30 | 0.194 | 0.302 | 0.409 | 0.7 | 0.3 | 1.29 | 1.4 | 32.33 | 18.03 | 0.07 | 4.57 | 9.2 |
| | 60 | 0.209 | 0.319 | 0.415 | 0.5 | 0.22 | 1.34 | 0.9 | 34.83 | 16.91 | 0.06 | 4.69 | 9.6 |
| | 100 | 0.208 | 0.318 | 0.414 | 0.38 | 0.22 | 1.4 | 0.52 | 34.68 | 16.41 | 0.05 | 4.82 | 9.6 |
| | 200 | 0.198 | 0.305 | 0.408 | 0.05 | 0.28 | 1.46 | 0.29 | 33.03 | 16.13 | 0.05 | 5 | 9.6 |
| NG04309279 / UBIAJA | 5 | 0.154 | 0.264 | 0.399 | 1 | 0.63 | 1.23 | 2.79 | 25.52 | 25.17 | 0.12 | 5.82 | 25.4 |
| | 15 | 0.165 | 0.277 | 0.402 | 0.85 | 0.51 | 1.25 | 2.37 | 27.49 | 24.26 | 0.09 | 5.9 | 22.3 |
| | 30 | 0.181 | 0.295 | 0.408 | 0.7 | 0.37 | 1.28 | 1.8 | 30.23 | 23.1 | 0.07 | 5.99 | 21.6 |
| | 60 | 0.196 | 0.311 | 0.414 | 0.5 | 0.28 | 1.33 | 1.16 | 32.71 | 21.97 | 0.06 | 6.1 | 22.6 |
| | 100 | 0.196 | 0.31 | 0.413 | 0.38 | 0.28 | 1.39 | 0.68 | 32.75 | 21.32 | 0.05 | 6.24 | 22.7 |
| | 200 | 0.186 | 0.297 | 0.408 | 0.05 | 0.35 | 1.45 | 0.37 | 30.96 | 20.88 | 0.05 | 6.42 | 22.7 |
| NG04340916 / IKENNE | 5 | 0.195 | 0.312 | 0.415 | 1 | 0.28 | 1.4 | 2.47 | 32.57 | 22.94 | 0.12 | 5.1 | 15.6 |
| | 15 | 0.208 | 0.326 | 0.42 | 0.85 | 0.22 | 1.42 | 2.09 | 34.67 | 22.25 | 0.09 | 5.17 | 13.6 |
| | 30 | 0.225 | 0.344 | 0.428 | 0.7 | 0.15 | 1.45 | 1.59 | 37.55 | 21.25 | 0.07 | 5.26 | 13.2 |
| | 60 | 0.239 | 0.359 | 0.436 | 0.5 | 0.11 | 1.5 | 1.02 | 39.97 | 20.14 | 0.06 | 5.38 | 13.8 |
| | 100 | 0.239 | 0.359 | 0.435 | 0.38 | 0.11 | 1.56 | 0.59 | 40.08 | 19.45 | 0.05 | 5.52 | 13.8 |
| | 200 | 0.228 | 0.346 | 0.428 | 0.05 | 0.14 | 1.62 | 0.33 | 38.08 | 18.86 | 0.05 | 5.7 | 13.8 |
| NG04283331 / AGO | 5 | 0.136 | 0.237 | 0.391 | 1 | 0.94 | 1.31 | 2.43 | 22.72 | 22.25 | 0.12 | 4.9 | 9.9 |
| | 15 | 0.148 | 0.25 | 0.394 | 0.85 | 0.75 | 1.33 | 2.07 | 24.69 | 21.45 | 0.09 | 4.97 | 8.6 |
| | 30 | 0.164 | 0.269 | 0.398 | 0.7 | 0.55 | 1.36 | 1.57 | 27.4 | 20.29 | 0.07 | 5.06 | 8.4 |
| | 60 | 0.179 | 0.285 | 0.403 | 0.5 | 0.42 | 1.41 | 1.01 | 29.78 | 19.06 | 0.06 | 5.18 | 8.8 |
| | 100 | 0.178 | 0.283 | 0.402 | 0.38 | 0.42 | 1.47 | 0.59 | 29.67 | 18.58 | 0.05 | 5.32 | 8.8 |
| | 200 | 0.167 | 0.268 | 0.397 | 0.05 | 0.54 | 1.52 | 0.33 | 27.79 | 18.32 | 0.05 | 5.5 | 8.8 |
| NG04261714 / IBADAN | 5 | 0.106 | 0.187 | 0.384 | 1 | 1.88 | 1.35 | 1.23 | 17.82 | 16.66 | 0.12 | 5.34 | 9.1 |
| | 15 | 0.116 | 0.2 | 0.385 | 0.85 | 1.55 | 1.37 | 1.04 | 19.56 | 16.1 | 0.09 | 5.41 | 8 |
| | 30 | 0.131 | 0.218 | 0.387 | 0.7 | 1.17 | 1.4 | 0.8 | 22.02 | 15.51 | 0.07 | 5.51 | 7.9 |
| | 60 | 0.146 | 0.235 | 0.389 | 0.5 | 0.89 | 1.45 | 0.51 | 24.37 | 14.59 | 0.06 | 5.62 | 8 |
| | 100 | 0.144 | 0.232 | 0.388 | 0 | 0.93 | 1.51 | 0.3 | 24.1 | 14.07 | 0 | 5.75 | 8.1 |
| | 200 | 0.134 | 0.218 | 0.386 | 0 | 1.16 | 1.57 | 0.19 | 22.46 | 13.61 | 0 | 5.94 | 8.1 |
| NG04171020 / MOKWA | 5 | 0.198 | 0.312 | 0.413 | 1 | 0.27 | 1.3 | 1.64 | 33.07 | 20.59 | 0.12 | 5.82 | 17.3 |
| | 15 | 0.21 | 0.325 | 0.419 | 0.85 | 0.21 | 1.32 | 1.39 | 35.11 | 19.91 | 0.09 | 5.89 | 15.1 |
| | 30 | 0.225 | 0.342 | 0.426 | 0.7 | 0.15 | 1.35 | 1.06 | 37.65 | 19 | 0.07 | 5.99 | 14.7 |
| | 60 | 0.239 | 0.357 | 0.433 | 0.5 | 0.11 | 1.41 | 0.68 | 39.98 | 17.99 | 0.06 | 6.1 | 15.4 |
| | 100 | 0.238 | 0.356 | 0.432 | 0.38 | 0.11 | 1.47 | 0.39 | 39.91 | 17.27 | 0.05 | 6.24 | 15.5 |
| | 200 | 0.229 | 0.344 | 0.426 | 0.05 | 0.14 | 1.52 | 0.21 | 38.32 | 16.87 | 0.05 | 6.41 | 15.5 |
| NG04184010 / ABUJA | 5 | 0.137 | 0.23 | 0.389 | 1 | 1.01 | 1.21 | 2.19 | 22.94 | 17.15 | 0.12 | 5.55 | 14.1 |
| | 15 | 0.149 | 0.243 | 0.391 | 0.85 | 0.81 | 1.22 | 1.86 | 24.85 | 16.45 | 0.09 | 5.61 | 12.3 |
| | 30 | 0.165 | 0.262 | 0.395 | 0.7 | 0.59 | 1.26 | 1.42 | 27.49 | 15.64 | 0.07 | 5.7 | 12 |
| | 60 | 0.179 | 0.279 | 0.4 | 0.5 | 0.43 | 1.31 | 0.91 | 29.95 | 14.8 | 0.06 | 5.82 | 12.4 |
| | 100 | 0.178 | 0.277 | 0.399 | 0 | 0.44 | 1.37 | 0.52 | 29.78 | 14.23 | 0 | 5.96 | 12.5 |
| | 200 | 0.169 | 0.264 | 0.395 | 0 | 0.55 | 1.42 | 0.31 | 28.18 | 13.73 | 0 | 6.14 | 12.5 |

SLLL: Lower limit of plant extractable soil water ($cm^3$)
SDUL: Drained upper limit ($cm^3$)
SSAT: Upper limit, saturated ($cm^3$)
SRGF: Soil Root growth factor
SSKS: Saturated hydraulic conductivity (cm/h)
SBDM: Bulk density ($g/cm^3$)
SLOC: Organic carbon (%)
SLCL: Clay (%)
SLSI: Silt (%)
SLNI: Total nitrogen (%)
SLHW: pH in water
SCEC: Cation exchange capacity (cmol/kg)

### Supplementary Methods

The Generalized Likelihood Uncertainty Estimation (GLUE) program as described by Beven & Binley (1992), He et al. (2010), and updated by He et al. (2021).

#### a. Develop prior parameter distributions

Random sampling, based on the prior distribution of parameters, is generally implemented in GLUE. The ranges of the parameters were defined and stored in a file formatted for the GLUE program under DSSAT 4.8.0.

#### b. Generate random parameter sets from the prior parameter distributions

The parameters sets were generated using a Monte Carlo sampling of distributions implemented in a MATLAB program. From the point of view of Monte Carlo sampling in the GLUE method, more parameter sets lead to more stable results (Li et al., 2018). Thus, a total of 10,000 (the minimum recommended is 3,000) random parameter sets were generated from the prior distribution.

#### c. Run the MANIHOT-Cassava model

The MANIHOT-Cassava model was run with each parameter set generated above by introducing the set into the standard genetic input file (cultivars file). For each run, the program identified all experiments in the DSSAT cassava data files involving the accession selected and these were used in the coefficient estimation process. The crop growth simulation outputs of dry aboveground biomass for each parameter set were tabulated for use in the GLUE likelihood calculations.

#### d. Calculate the likelihood values and probability

The observed data on dry aboveground biomass were used along with the corresponding simulated outputs to compute the likelihood value. The Likelihood Function is the product of these individual likelihood values. The Gaussian likelihood function used in this study was recommended by He *et al.* (2010):

$$L[\theta_i|O] = \Pi_{j=1}^{M} \frac{1}{\sqrt{2\pi\sigma_o^2}} \exp[\frac{(O_j - P_j(\theta_i))^2}{2\sigma_o^2}] \qquad \text{Eq. 4}$$

$L[\theta_i|O]$ = likelihood value of parameter set i, given the set of observations
$\theta_i$ = i$^{th}$ parameter set (I = 1, 2, 3, ...N). N = 10,000 in our case.
$O_j$ = j$^{th}$ observation
$P_j(\theta_i)$ = model output for the j$^{th}$ environment under parameter set $\theta_i$
$\sigma_o^2$ = variance of model error
M = number of observations.
The posterior probability $p(\theta_i)$, of each parameter set $\theta_i$ is computed with the Bayesian equation as follows:

$$p(\theta_i) = \frac{L(\theta i|O)}{\sum_{i=1}^{N} L(\theta i|O)} \qquad \text{Eq. 5}$$

**e. Construct posterior distribution and statistics**

The pair of parameter sets and probabilities [$\theta_i$, $p(\theta_i)$] were used to compute the mean, and the variance of each parameter $\theta_i$:

$$u_{post}(\theta) = \sum_{i=1}^{N} p(\theta_i) \,.\, \theta_i \qquad \text{Eq. 6}$$

$$\sigma_{post}^2(\theta) = \sum_{i=1}^{N} p(\theta_i) \,.\, \left(\theta_i - u_{post}(\theta)\right)^2 \qquad \text{Eq. 7}$$

$u_{post}(\theta_i)$ = mean of the posterior distribution
$\sigma_{post}^2$ = variance of the posterior distribution
